# Genome modularization reveals overlapped gene topology is necessary for efficient viral reproduction

**DOI:** 10.1101/2020.06.10.143693

**Authors:** Bradley W Wright, Juanfang Ruan, Mark P Molloy, Paul R Jaschke

**Author notes:** Corresponding Author Addresses: Paul R. Jaschke, Department of Molecular Sciences, Macquarie University, Sydney, NSW 2109, AUSTRALIA, Mark P. Molloy, Faculty of Medicine and Health, Northern Clinical School, The University of Sydney, Sydney, NSW 2006, AUSTRALIA.

## Abstract

Sequence overlap between two genes is common across all genomes, with viruses having high proportions of these gene overlaps. The biological function and fitness effects of gene overlaps are not fully understood, and their effects on gene cluster and genome-level refactoring are unknown. The bacteriophage ϕX174 genome has ∼26% of nucleotides involved in encoding more than one gene. In this study we use an engineered ϕX174 phage containing a genome with all gene overlaps removed, to show that gene overlap is critical to maintaining optimal viral fecundity. Through detailed phenotypic measurements we reveal that genome modularization in ϕX174 causes virion replication, stability, and attachment deficiencies. Quantitation of the complete phage proteome across an infection cycle reveals almost half the proteins display abnormal expression patterns. Taken together, we have for the first time comprehensively demonstrated that gene modularization severely perturbs the coordinated functioning of a bacteriophage replication cycle. This work highlights the biological importance of gene overlap in natural genomes and that reducing gene overlap disruption should be an integral part of future genome engineering projects.

## INTRODUCTION

The arrangement of genes within a genome is well recognised to be an important feature in the regulation of gene expression (Cavalli & Misteli, 2013, Chen & Stein, 2006, Harmston, Ing-Simmons et al., 2017, Lanctôt, Cheutin et al., 2007, Lawrence, 2002, Lim, Lee et al., 2011, Tsochatzidou, Malliarou et al., 2017) and as such, gene order tends to be evolutionarily conserved (Dandekar, Snel et al., 1998, Davila Lopez, Martinez Guerra et al., 2010, Herniou, Olszewski et al., 2003, Minot, Wu et al., 2012, Poyatos & Hurst, 2007, Singer, Lloyd et al., 2004, Tamames, 2001). Studies have investigated implications of ectopic gene expression revealing functional significance to gene order (Endy, You et al., 2000, Flanagan, Schoeb et al., 2003, Pesko, Voigt et al., 2015, Wakimoto & Hearn, 1990). For example, studies in bacteriophage T7 have highlighted the criticality of gene ordering in achieving the upper limits of fitness (Endy et al., 2000, Springman, Badgett et al., 2005), and significantly, this fitness attenuating gene re-ordering is difficult to purge even after long-adaptation cycles (Cecchini, Schmerer et al., 2013). Therefore, despite important functional significance revealed by these studies, rules surrounding the relationship of genome architecture and phenotypic outcome are still only loosely defined.

In addition to gene order, another related feature of genome topology has been gene sequence overlap, which is common across viral (Belshaw, Pybus et al., 2007, Brandes & Linial, 2016), prokaryotic (Huvet & Stumpf, 2014, Johnson & Chisholm, 2004) and eukaryotic (Chung, Wadhawan et al., 2007, Mouilleron, Delcourt et al., 2015, Neme & Tautz, 2013) genomes, and as such, is increasingly thought to be a critical component of genome architecture (Johnson & Chisholm, 2004, Pavesi, Vianelli et al., 2018, Sanna, Li et al., 2008, Schlub & Holmes, 2020, Veeramachaneni, Makalowski et al., 2004). In the model eukaryote yeast, overlaps are prevalent, with approximately 18% of genes overlapping, majority of which lie in untranslated regions (UTR) (Sun, Yang et al., 2015). In viruses, gene overlaps are more prevalent and seem to be a result of hard constraints put on genome size as a result of capsid assembly and stability, restricting genome length and nucleotide composition (Brandes & Linial, 2016, Chirico, Vianelli et al., 2010). Once in place, gene overlaps also seem to have some role in gene expression regulation (David, 2000, Jin, Vacic et al., 2008, Johnson & Chisholm, 2004, Lapidot & Pilpel, 2006).

In prokaryotes and viruses, whereby gene overlap is often found within open reading frames (Johnson & Chisholm, 2004, Schlub & Holmes, 2020), the overlap is thought to arise from mutations resulting in start codon creation within existing genes, or between adjacent genes by mutations removing start or stop codons (Delaye, DeLuna et al., 2008, Van Oss & Carvunis, 2019). This results in 5’ or 3’ extensions of the existing gene into the adjacent gene (Delaye et al., 2008, Van Oss & Carvunis, 2019). This phenomenon is often called overprinting (Keese & Gibbs, 1992) and may also occur on the anti-sense strand, typically resulting in the generation of non-protein encoding but functionally relevant cis or trans-acting RNA (Brantl, 2002).

Only a few studies have investigated changes to fitness from uncoupling (modularising) gene overlap (Chan, Kosuri et al., 2005, Fernandes, Faust et al., 2016, Ghosh, Kohli et al., 2012, Jaschke, Lieberman et al., 2012, Li, Ma et al., 2007). With one exception, in which the desired pathway product was increased (Li et al., 2007), modularisation has resulted in fitness loss (Chan et al., 2005, Fernandes et al., 2016, Ghosh et al., 2012, Jaschke et al., 2012), but to date, no study has defined the molecular mechanisms underlying these fitness defects.

The functional effects of gene overlap disruption are extremely pertinent to genome engineering and refactoring projects, which frequently disrupt sequence overlaps during reconfiguration of native genes and pathways. A number of projects have cautiously sought to minimise the potential impacts of overlap disruption through explicit design choices such as overlap sequence duplication (Baba, Ara et al., 2006, Hutchison, Chuang et al., 2016, Lajoie, Kosuri et al., 2013), but these efforts to address the biological implications of gene overlap disruption have been ad-hoc. As such, there is no consensus regarding how to treat gene overlaps in genome engineering and refactoring projects.

In this study, we comprehensively define the range of phenotypes displayed by a whole genome modularised bacteriophage that was constructed previously (Jaschke et al., 2012). The aptly named ‘decompressed’ ϕX174 had all instances of gene overlap modularised, which had previously constituted 16.8% of the wild-type genome. The decompressed ϕX174 genome modifications removed gene overlaps of both entirely overprinted genes (genes B within A, K within A and C, and E within D) as well as start/stop overlaps of tandem same-strand gene overlaps (genes B-K, A-C, K-C, C-D, D-J) (Figure 1), while maintaining absolute genome length, codon usage bias, and G+C content (Jaschke et al., 2012).

**Figure 1:**
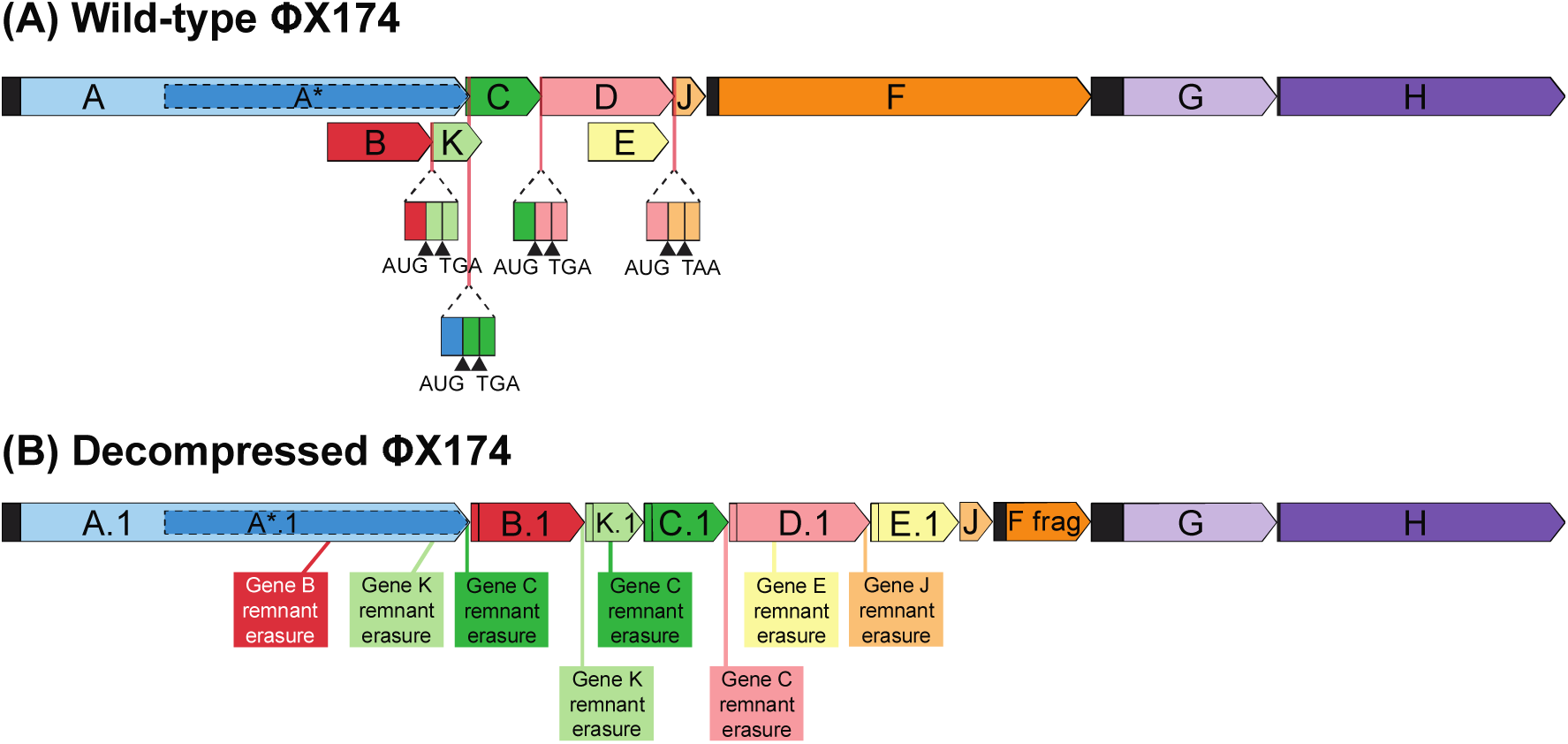
Wild-type and decompressed ϕX174 genome topologies. (A) In the wild-type ϕX174 genome, genes B and E are examples of overprinted gene overlap while tandem same-strand start/stop codon overlaps are shown for overlaps between genes B-K, A-C, C-D, and D-J. (B) In the decompressed ϕX174 genome all overprinted genes and start/stop codon overlaps are removed. Locations of synonymous codon modification to remove legacy start codons and ribosome binding sites are shown. Genes with the .1 nomenclature have had synonymous codon changes in at least one location within the gene.

Initial characterization of the resulting phage revealed comparable lysis efficiency with respect to the wild-type strain under the limited conditions tested (Jaschke et al., 2012). To build upon that work, here we have deeply characterized decompressed ϕX174 using targeted proteomics to temporally resolve the complete proteome of decompressed and wild-type ϕX174 across an entire infection cycle. We find that decompressed ϕX174 conceals severe deficiencies in several key replication proteins and displays a broad range of phenotypic issues. As a result of this work we gain new insights into the effect of genome modularisation and highlight the need to consider gene overlaps as a critical component of genome architecture, especially in the context of future genome engineering projects.

## RESULTS AND DISCUSSION

### Decompressed ϕX174 displays comparable lysis timing but impaired progeny production

To understand more deeply the phenotypic effects of removing all overlaps from the genome of bacteriophage ϕX174, we first performed plaque assays and liquid growth assays of decompressed ϕX174. Double-agar overlay plaque assays of wild-type and decompressed ϕX174 phage preparations infecting *Escherichia coli* C122(pJ804(Gene F)) strain under gene F inducing conditions (necessary for decompressed ϕX174 reproduction) showed that the decompressed ϕX174 plaque size was significantly smaller at 4.1 ± 0.9 mm^2^ compared to 29.2 ± 8.0 mm^2^ for wild-type ϕX174 (p-value <0.00001) (Figure 2A).

**Figure 2:**
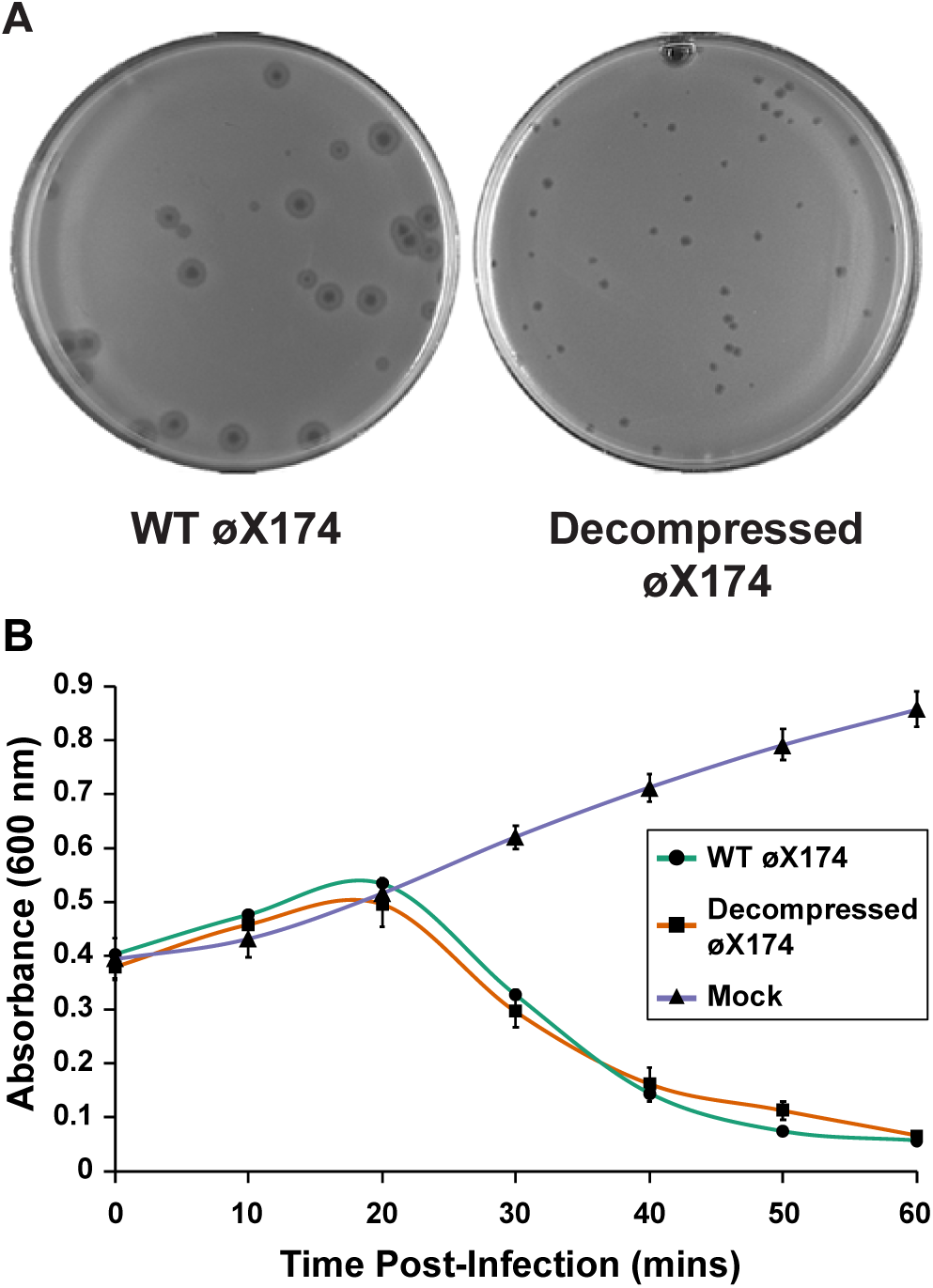
Decompressed ϕX174 lyses host at similar efficiency to wild-type ϕX174 at high multiplicity of infections but has reduced fecundity overall. (A) Double-agar overlay plates of wild-type ϕX174 and decompressed ϕX174 infecting host *E. coli* C122(pJ804(Gene F)). Wild-type ϕX174 plates contained 2 mM glucose and decompressed ϕX174 plates contained 2 mM rhamnose. Plate diameter = 85 mm. (B) Lysis curve at multiplicity of infection (MOI) = 5. Error bars represent one standard deviation (n = 2-3).

During phage infection, there is an intricate balance between progeny production and lysis initiation, and there are a range of factors that can decrease plaque size, including decreased latent period or burst-size (Abedon & Culler, 2007a, Abedon & Culler, 2007b, Gallet, Kannoly et al., 2011). The ϕX174 protein E mediates host lysis, with an estimated 500 molecules sufficient to cause membrane disruption and cell death (Zheng, Struck et al., 2009). If decompressed ϕX174 was producing protein E sooner in the infection cycle than wild-type ϕX174 or was producing less E protein, that may explain the reason for smaller plaque size due to reduced progeny production from premature lysis or inefficient lysis.

To determine whether lysis-timing or lack of effective lysis was the cause of the decompressed ϕX174 small-plaque phenotype, we infected exponentially growing *E. coli* C122(pJ804(Gene F)) strain with either decompressed or wild-type ϕX174 with a multiplicity of infection (MOI) = 5 and measured culture absorbance for 60-minutes. We used a high phage load to ensure the majority (>99%) of host cells had one or more phage attached and we were only observing the effects of one infection cycle. The results of this time-course showed that decompressed and wild-type ϕX174 have comparable lysis timing under our conditions (Figure 2B) with no evidence for phage cross-contamination (Appendix Figure S1).

We next measured the burst-size of decompressed ϕX174 to determine if the small-plaque phenotype could be due to fewer viable progeny produced per infected cell. We determined the burst-size of wild-type ϕX174 to be 145 ± 66 phage produced per cell while decompressed ϕX174 produced only 16 ± 2 phage per cell. Previous measurements of wild-type ϕX174 burst-size have reported between 160-180 phage produced per cell (Brown, Stancik et al., 2013, Gillam, Atkinson et al., 1985, Hutchison & Sinsheimer, 1966).

The significant burst-size deficiency in decompressed ϕX174 is the likely reason for the small plaque size in this engineered phage (Figure 2A). Poor lysis is not the reason for altered burst-size in decompressed ϕX174 as lysis timing is not altered in a single-step growth experiment (Figure 2B). Additionally, we removed all unattached phage before infection initiation in this assay, effectively excluding aberrant host attachment or recognition as the cause of this burst-size deficiency. We next assessed whether phage virion structure or genome ejection into cells could be the cause of the reduced burst-size. The decompressed ϕX174 genome encodes proteins with the same amino acid sequence as the wild-type ϕX174 genome, as such, we expect any structural changes to be either from disruption of the intimate genome-capsid interactions due to packaging a genome of different sequence or from inefficient assembly due to altered protein levels during capsid assembly.

### Decompressed ϕX174 virion has poor heat stability and attachment efficiency

To identify whether decompressed ϕX174 virions are less stable than wild-type due to the differences in the packaged genome, we measured the effect of heat stress on phage infectivity. Samples of purified wild-type and decompressed ϕX174 virions were subjected to 60°C over a range of times and the remaining infectivity was compared to untreated samples. Decompressed ϕX174 virions were rapidly inactivated by the heat treatment after only 5-minutes, while wild-type ϕX174 was slightly more infectious after this treatment (Figure 3A). A longer 15-minute treatment also significantly decreased decompressed ϕX174 infectivity more than wild-type (Figure 3A).

**Figure 3:**
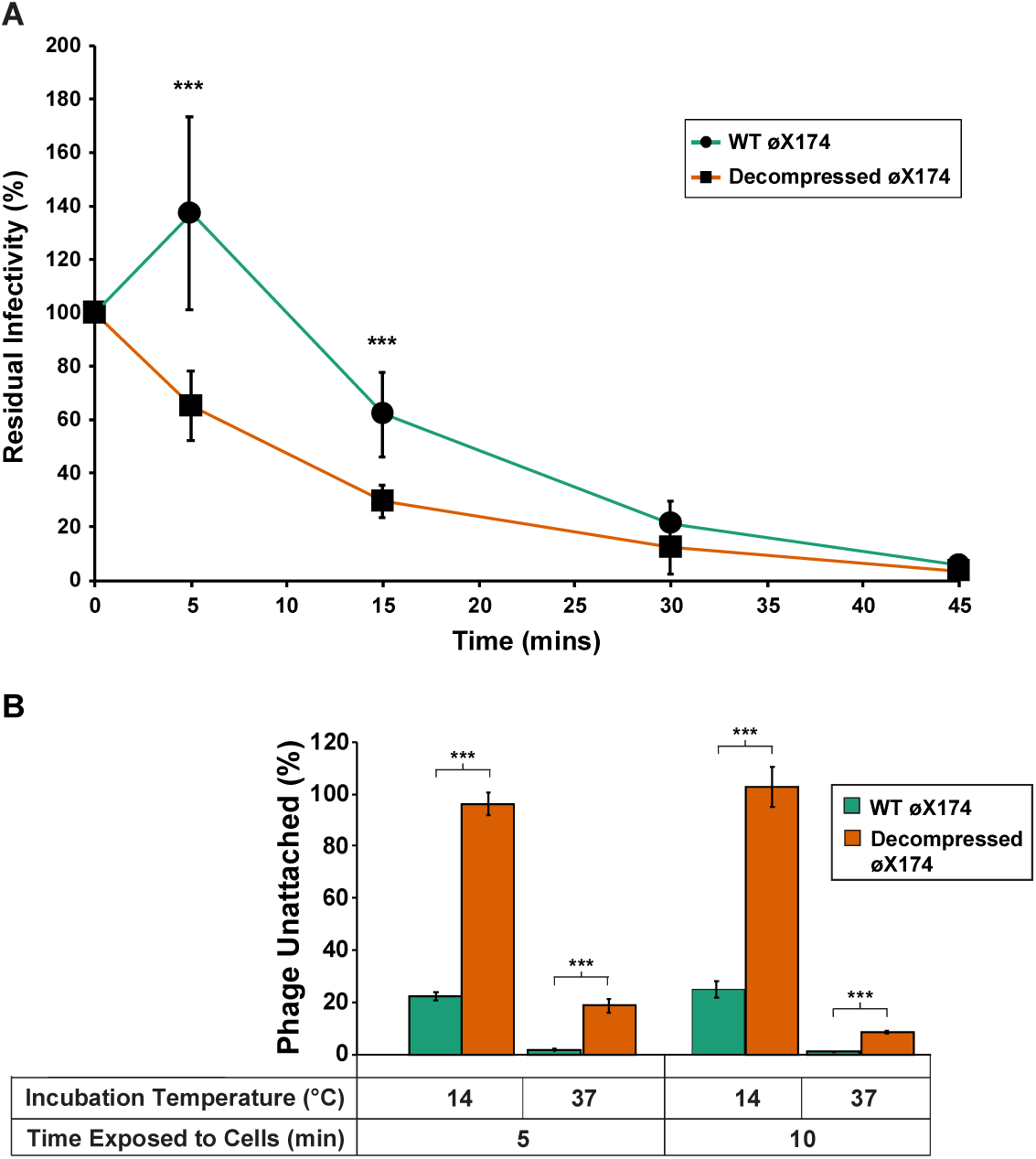
Decompressed ϕX174 is both less heat stable and poorer at attaching to host cells than wild-type ϕX174. (A) Heat stability of phage. Samples of decompressed ϕX174 and wild-type ϕX174 virus were exposed to heat stress (60 °C) for 5, 15, 30, and 45-minutes followed by measurement of residual infectivity by double-agar plate plaque enumeration. Error bars represent one standard deviation (n = 9). *** p-value < 1.0 × 10^−4^. (B) Viral attachment assay. Lysis resistant *E. coli* C900 Δ*slyD* cells were incubated at the specified temperature with either wild-type ϕX174 or decompressed ϕX174 phage for either 5 or 10-minutes followed by enumerating unattached phage using the double-agar overlay method. Error bars represent one standard deviation (n = 4). *** p-value < 1.0 x 10^−4^.

This result indicates that the modularised genome of decompressed ϕX174 is having a negative effect on the thermal stability of the virion. This is not totally unexpected, as studies addressing heat stability of viral capsids with genomic variation have revealed different thermal stability between counterparts (Emerson, Arankalle et al., 2005, Ojosnegros, García-Arriaza et al., 2011, Saha, Wong et al., 2014). The physical properties of constituents within capsid shells (for example, salt ions and nucleotides) have been shown to directly influence a capsid’s internal osmotic pressure, and consequently, mechanical stability (Evilevitch, Fang et al., 2008, Evilevitch, Lavelle et al., 2003, Ivanovska, Wuite et al., 2007, Roos, Bruinsma et al., 2010). So, while the decompressed ϕX174 genome encodes for the same genes and proteins, and is of the same absolute nucleotide length as the wild-type, it is reasonable to suggest that genome organisation is critical to maintaining thermal stability by maintaining optimal internal pressure and internal capsid interactions. Genome modularisation and the resulting alterations to the genome secondary structure (Appendix Figure S2) could alter the electrostatic interactions within the capsid (Belyi & Muthukumar, 2006, Erdemci-Tandogan, Wagner et al., 2014, Šiber, Božic et al., 2012) therefore directly influencing capsid rigidity. These complex interactions and their impact on particle stability and genome packaging, have been observed before in other viruses (Bauer, Li et al., 2015, Mateo, Luna et al., 2008, Ojosnegros et al., 2011, Snijder, Uetrecht et al., 2013).

Given the lower thermal stability of the decompressed ϕX174 virion we next measured the attachment efficiency to determine virion host-recognition and subsequent attachment. Attachment for wild-type and decompressed ϕX174 was measured at 14 °C and 37 °C. At 14 °C in starvation buffer ϕX174 adsorbs to the host cell without undergoing eclipse (irreversible structural change) (Denhardt & Sinsheimer, 1965, Newbold & Sinsheimer, 1970c). This characteristic of ϕX174 is particularly advantageous as it allows for the synchronisation of infection. In contrast, at 37 °C in starvation buffer ϕX174 undergoes an irreversible structural change leading to the partial ejection of the genome (Newbold & Sinsheimer, 1970c).

The results of the attachment assay at 14 °C showed that decompressed ϕX174 virions could not remain stably attached to host cells, as opposed to wild-type ϕX174 that had 77.8% (± 1.6%) virions remaining attached after incubation with hosts for 5 minutes (Figure 3B). At 37 °C decompressed ϕX174 virions attach more stably to host cells, with 81.3% (± 2.7%) left attached, but was still significantly less than wild-type under the same conditions (91.5% (±0.6 %), p-value = 0.00004). Longer 10-min incubations did not substantially improve relative decompressed ϕX174 attachment (Figure 3B). These attachment assay results suggest that decompressed ϕX174 virions can recognise the host cell receptor, as they are able to remain stably attached at 37°C and eclipse, but at 14°C, pre-eclipsed decompressed ϕX174 does not remain stably adsorbed to the host.

Together, these results provide evidence that the packaged genome of decompressed ϕX174 is impacting the properties of the virion, which in turn impairs attachment. Zeng, et al. (2017), measured the mechano-chemical properties of adeno-associated virus 2 and brome mosaic virus with different genomic cargo. They found that adding exogenous genetic material to these viral genomes has an influence on virus-substrate adhesion interactions with differing deformation and strong orientation biases for adsorption depending upon the genomic content (Zeng, Moller-Tank et al., 2017b). Further work by this group (Zeng, Hernando-Perez et al., 2017a) has illuminated the capsid orientation bias and deformity arising from adsorption in the structurally similar brome mosaic virus. The orientation bias as a result of the deformity of the capsid to maximise contact area would involve compressing the capsid inner surface and extending the outer surface (Zeng et al., 2017a), actions that will be impacted by properties of the encapsulated genome (Zeng et al., 2017b). As reflected by the attachment assay results, these mechano-chemical properties are likely perturbed in the decompressed ϕX174 virion. To further support the notion that the genome-capsid interactions have profound effects on the dynamic function of virions, previous work in ϕX174 has shown that altering the internal capsid environment through mutant J proteins also alters attachment efficiency (Bernal, Hafenstein et al., 2004, Hafenstein & Fane, 2002, Hafenstein, Chen et al., 2004). Additionally, altering the packaged genome sequence was also found to have profound effects on capsid biophysical properties (Hafenstein & Fane, 2002). These studies, along with our experiments, suggest that the genome of the decompressed ϕX174 virion is influencing the structural integrity and surface properties of the capsid, leading to perturbation of pre-eclipse attachment and capsid heat stability. To our knowledge, this study demonstrates for the first time that native gene architecture is critical for optimal capsid function.

### Decompressed ϕX174 capsid morphology is identical to wild-type

To identify any major structural aberrations in the decompressed ϕX174 capsid that may be causing the decreased attachment efficiency, we incubated purified virions with either DNase I or proteinase K and measured changes to viability. The results of this assay revealed that decompressed ϕX174 virions are slightly more stable under protease stress than wild-type (p-value 0.0173), but there was no significant difference under DNase I treatment, thereby making altered capsid permeability or partial genome ejection unlikely (Appendix Figure S3).

To characterize subtler variations in capsid structure between wild-type and decompressed ϕX174 we used cesium chloride (CsCl) gradient purification to isolate virions and examined them by transmission electron microscopy (TEM). Visualisation and subsequent 2D-classification of wild-type and decompressed ϕX174 virions revealed no observable capsid morphology differences (Figure 4). Additionally, the diameter of both decompressed and wild-type virions was measured to be 33 nm, in close agreement with wild-type measurements in other studies (McKenna, Xia et al., 1992, Sun, Roznowski et al., 2017, Yazaki, 1981). As no gross structural deformities were observed with the decompressed virion, we hypothesize that the mechanical properties of the capsid have been altered due to the packaging of the decompressed genome, thereby producing altered thermal stability and attachment issues (Figure 3).

**Figure 4:**
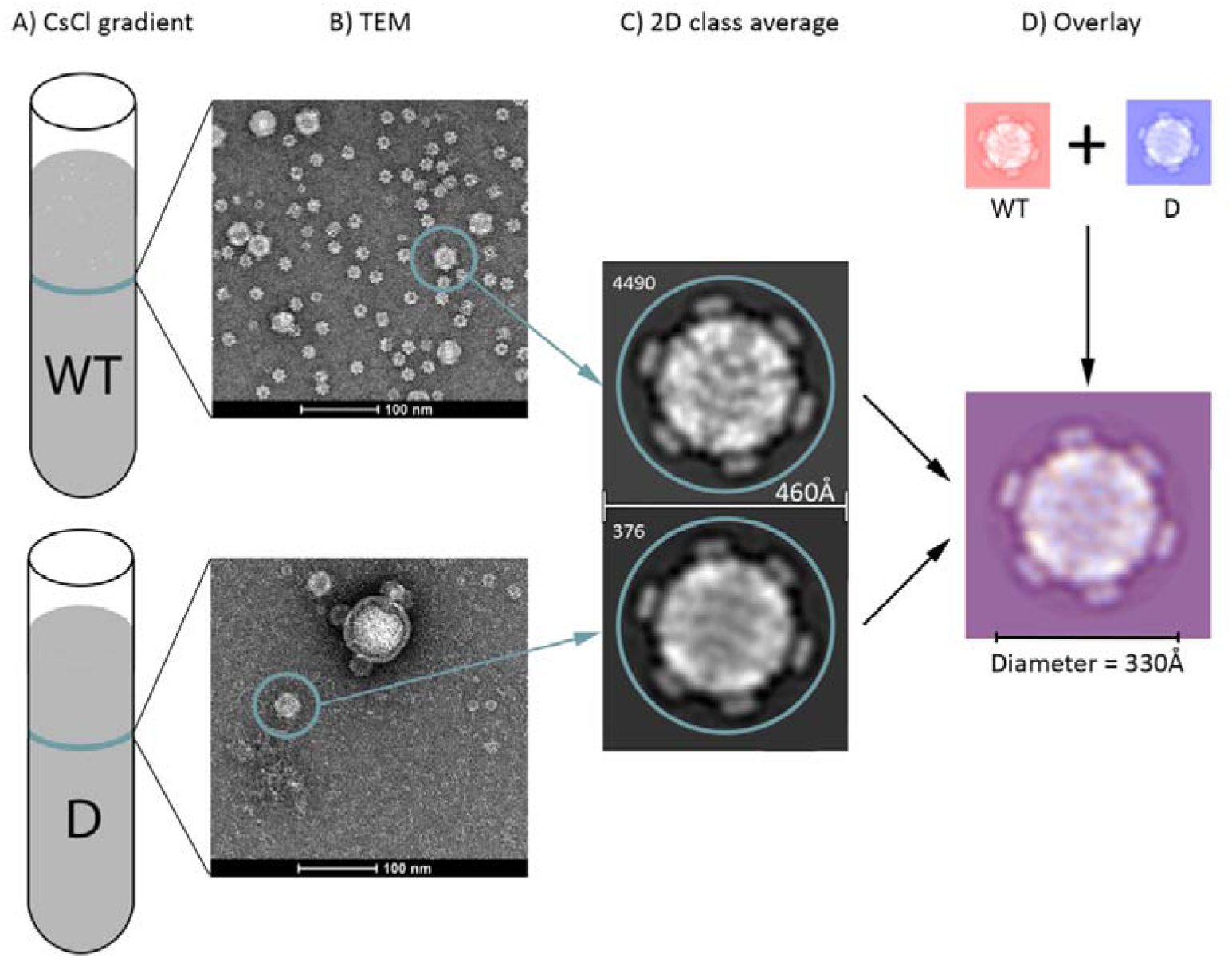
Decompressed ϕX174 and wild-type ϕX174 virion morphologies are indistinguishable. (A) Phage band isolated after CsCl step-gradient ultracentrifucation. WT = wild-type and D = Decompressed ϕX174. (B) Uranyl-acetate negative stained particles visualised using transmission electron microscopy (TEM). Phage particles found in both preparations (teal circle). (C) 2D class averaging of phage capsids highlighted in (B). 4490 particles averaged for the wild-type capsid, and 376 particles averaged for the decompressed ϕX174. Additional classifications and particle information available as Appendix Figures S4 and S5. (D) 2D class averaged images of capsids overlayed reveal no obvious morphology differences between wild-type ϕX174 and decompressed ϕX174.

Curiously, we also observed *E. coli* chaperone complex GroEL structures mixed with ϕX174-containing CsCl bands (Appendix Figure S4). As many viruses rely on host cellular chaperones to mediate their protein folding (Geller, Taguwa et al., 2012, Mayer, 2005, Xiao, Wong et al., 2010), including GroEL (Bouvaine, Boonham et al., 2011, Ding, Duda et al., 1995, Hanninen, Bamford et al., 1997), the co-purification of this large protein complex is not surprising, but has never been associated with ϕX174-infections before.

### Targeted proteomics reveals decompressed ϕX174 infections initiate protein production earlier than wild-type

To identify whether there are any defects in phage protein production that could explain the decompressed ϕX174 reduced burst-size, we quantified the level of individual ϕX174 proteins produced during infection of the *E. coli* host. We infected *E. coli* C122(pJ804(Gene F)) with either wild-type or decompressed ϕX174 phage at MOI = 1.2 and captured samples over a time-course. Proteins from four post-infection time-points (0, 15, 25, and 35-minutes) were extracted and analysed by parallel reaction monitoring mass spectrometry (PRM) (Peterson, Russell et al., 2012). PRM is a targeted mass spectrometry acquisition method that enables highly sensitive and reproducible measurements of pre-determined peptide ions (Peterson et al., 2012), and was therefore a well-suited analytical method for quantitation of phage proteins *in vivo*. This approach successfully detected and quantified the relative abundance of phage proteins from within the dynamic background of the *E. coli* proteome, enabling comparative analyses of the wild-type and decompressed ϕX174 phage proteomes across the infection cycle. For quantitation, each phage protein was measured using at least two peptides (except for proteins K, J, and E – where only one tryptic peptide was suitable (Appendix Dataset S1)).

Using PRM we observed viral protein expression in the decompressed ϕX174 samples at time point 0-minutes (Figure 5 and Appendix Figures S6 and S7). Immediate expression was observed for all proteins except A, B, C, and K. The F protein was constitutively expressed from plasmid pJ804(Gene F). In contrast, proteins produced from wild-type ϕX174 were not observed until at least 15-minutes post-infection (Figure 5). This surprising result is in contrast to previous literature that showed infection synchronisation conditions (incubation with cells at <19°C and in starvation buffer) completely inhibited wild-type ϕX174 eclipse and DNA ejection phases (Denhardt & Sinsheimer, 1965, Newbold & Sinsheimer, 1970b, Newbold & Sinsheimer, 1970c). Therefore, this could indicate that decompressed ϕX174 phage possessed some altered infection dynamics under the cold starvation conditions (Material and Methods). Temperature sensitive mutants of ϕX174 have been shown to exhibit altered eclipse and DNA injection characteristics (Dalgarno & Sinsheimer, 1968, Dowell, 1967, Ilag & Incardona, 1993, Incardona, 1974), possibly as a result of structural changes (Dalgarno & Sinsheimer, 1968, Dowell, 1967, Ilag & Incardona, 1993), and may be evidenced by altered thermal stability of the phage virions (Dalgarno & Sinsheimer, 1968, Dowell, 1967). The viral protein present at time point 0-minutes (Figure 5) and lower heat stability (Figure 3A) together show altered properties of the decompressed ϕX174 capsid, likely as a result of the decompressed genome packaged therein.

**Figure 5:**
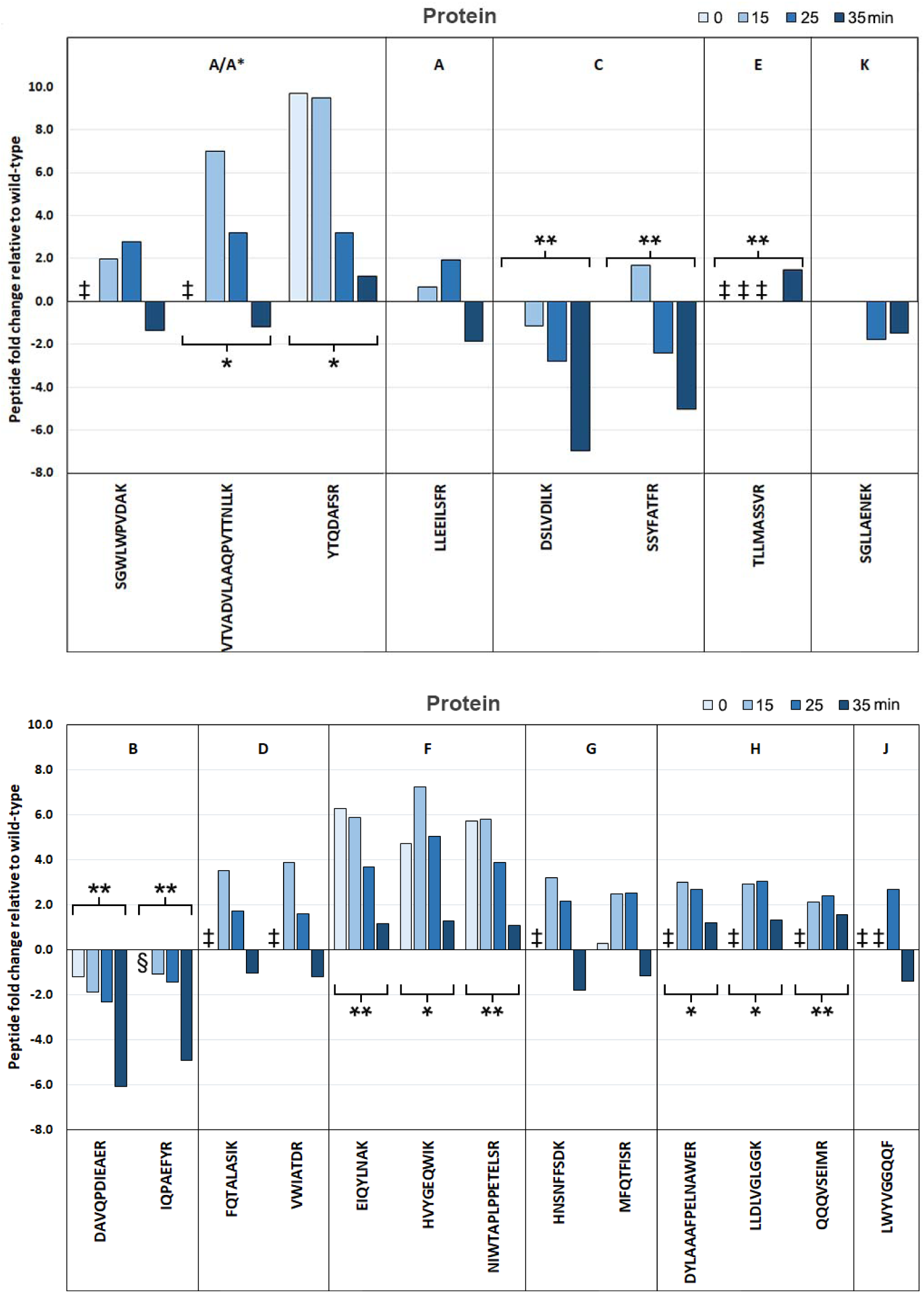
Mass spectrometric quantification of ϕX174 proteins produced during infection shows altered expression levels of proteins involved in genome replication and capsid packaging and assembly in decompressed ϕX174. PRM quantification of targeted viral peptides across four time-points comparing decompressed relative to wild-type. (Top) Fold-change of peptides from replication associated proteins. (Bottom) Fold-change of peptides from capsid associated proteins. ‡ = Peptide detected in decompressed ϕX174 but not in the wild-type ϕX174, § = Peptide detected in wild-type ϕX174 but not in the decompressed ϕX174. Two-factor ANOVA significance * p-value <0.05. ** p-value <0.005.

### Proteomics reveals decompressed ϕX174 infection is deficient in proteins B and C, whilst protein A* is overexpressed

Next we determined whether any ϕX174 proteins (Appendix Table 1) produced during the infection cycle were altered between wild-type and decompressed ϕX174. Two-factor ANOVA analysis was performed on each peptide with independent variables set to time and phage strain. The results of this analysis showed there were significant alterations in the expression of proteins A*, B, C, E, F, and H during decompressed ϕX174 infection (Figure 5, Appendix Datasets S1 and S2).

The lysis protein E was produced at high levels in decompressed ϕX174 across all time points relative to the wild-type (Figure 5). In contrast, in wild-type ϕX174, protein E was only detectable at 35-minutes post-infection (Appendix Figure S6). This result is unexpected, as we measured lysis time of decompressed ϕX174 as comparable to that of wild-type at MOI=5 (Figure 2B) and therefore expected this protein’s production to be similar between the strains in the proteomics experiment. Premature lysis or damage to the host cell as a result of expression of the lysis protein would most certainly impact viral infection fidelity as progeny production would be interrupted. Constant protein E expression from decompressed ϕX174 during infection indicates that the modularisation of the genome may have modified the lysis protein E regulation from wild-type, resulting in leaky expression. The gene for protein E was originally completely encoded within gene D, and thus modularisation has resulted in the E gene being moved 296 nucleotides downstream from its original site within ϕX174 transcripts (Figure 6).

**Figure 6:**
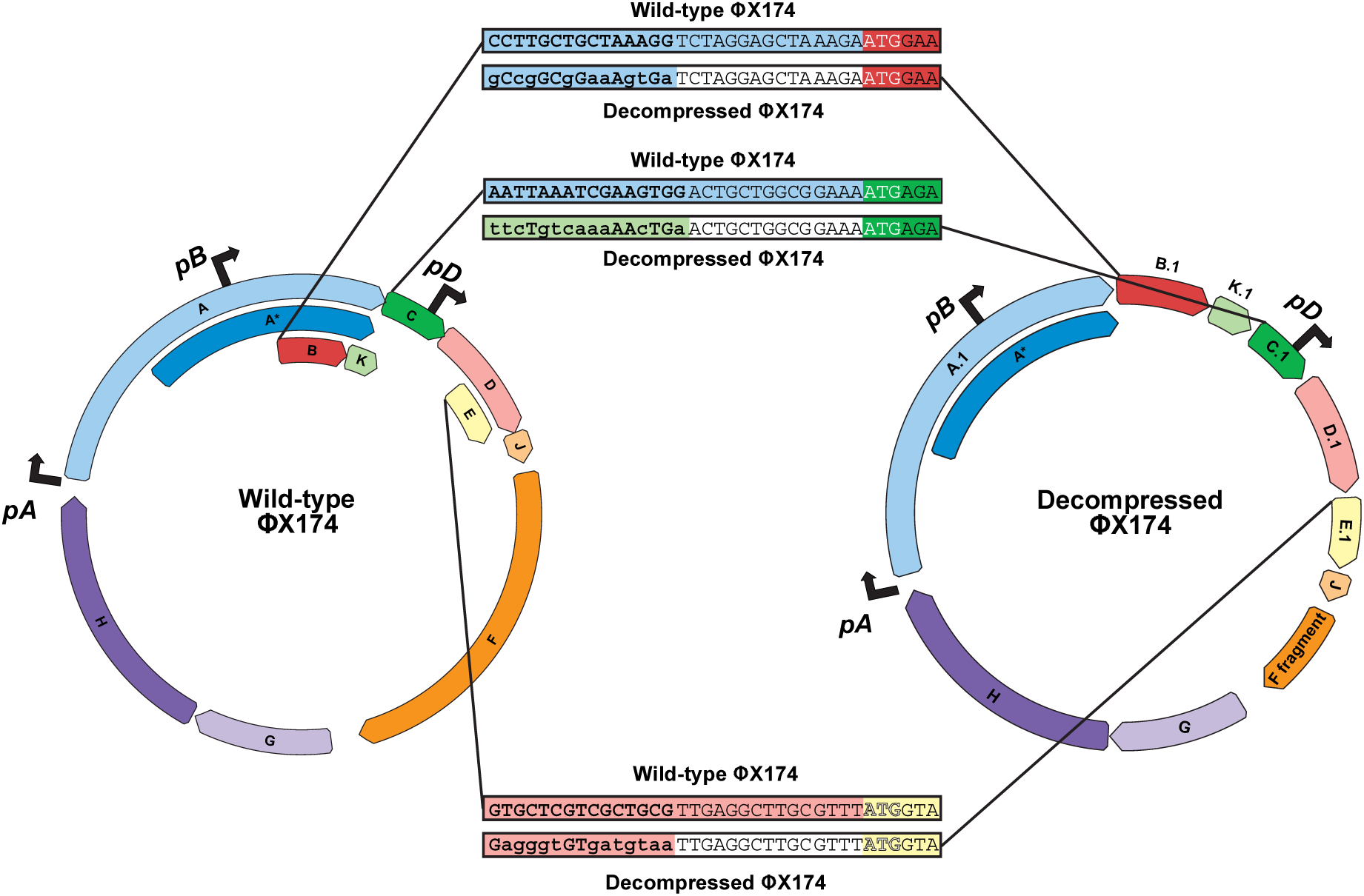
The heavily modified 5’-UTR of genes B, C, and E in the decompressed ϕX174 genome may result in altered protein expression. Gene B, C, and E sequences in the wild-type and decompressed ϕX174 genomes are highlighted (boxed) and show the region 5’ of the start codons (white bolded ATG text). Colored background to text indicates which gene the sequence belongs to: gene A (light blue), gene K (light green), gene C (dark green), gene D (pink), gene E (yellow), no gene (white). Wild-type gene C ATG start codon overlaps with gene A TGA stop codon. Bolded text and lower case nucleotides show positions in the 5’-UTRs of decompressed ϕX174 genes that differ from wild-type sequence.

In contrast to protein E, we found protein C expression was significantly diminished in the decompressed ϕX174 infection (Figure 5). In the wild-type genome, gene C has significant levels of upstream sequence overlap with gene K as well as an overlapping start-stop codon junction with genes A/A*\** (Figure 6). The modularisation of these junctions, resulting in a modified 5’-UTR region and removal of the A/C overlap may be responsible for decreased protein C expression.

During ϕX174 infections, protein C is responsible for the shift from single-strand genome production seeding the generation of more dsDNA replicative forms (stage II) to the production of ssDNA genomes destined for packaging within capsids to assemble infectious virions (stage III) (Aoyama & Hayashi, 1986, Doore, Baird et al., 2014). More specifically, protein C functions by competitively binding to the ϕX174 origin of replication with antagonistic partner, host protein single-strand binding protein (SSB) (Doore et al., 2014). If protein C binds, then a round of stage III DNA synthesis occurs in which the production of single-stranded genomic DNA occurs, followed by packaging into the viral procapsid (Aoyama & Hayashi, 1986). If SSB protein binds, then stage II synthesis continues (Aoyama & Hayashi, 1986). It is known that an increase in host SSB protein relative to protein C during stationary phase infection results in reduced plaque sizes (Doore et al., 2014). The ∼6-fold decreased expression of protein C at 35-minutes post-infection likely results in a reduced ability to transition from stage II to stage III DNA synthesis and less efficient packaging of ssDNA genomes into available procapsids. Our observation of reduced protein C may also explain the reduced burst and plaque sizes for decompressed ϕX174 (Figure 2A).

We also found that protein A* was significantly upregulated during decompressed ϕX174 infections. Although protein A and A* share 100% sequence identity, we were able to discriminate between protein A and A* production through the measurement of the unique A peptide, LLEEILSFR. By comparing LLEEILSFR levels to three other peptides shared by proteins A and A* we observed that protein A* is highly upregulated while protein A is not (Figure 5).

Protein A* controls the delicate balance between packaging and replication during ϕX174 infections (Roznowski, Doore et al., 2019). This function is accomplished through binding within the J-F intergenic region of the ϕX174 genome. This A* binding leads to inhibiting DNA unwinding in this region and disruption of protein A rolling circle genome amplification (Eisenberg & Ascarelli, 1981, Roznowski et al., 2019, van Mansfeld, van Teeffelen et al., 1986). Therefore stochiometry differences between proteins A and A* brought about by A* upregulation in decompressed ϕX174, are likely to cause issues with genome replication and the maintenance of genome packaging fidelity.

Protein B is an essential internal scaffolding protein that is needed for the assembly of the procapsid, an intermediary capsid particle devoid of the phage genome (Dokland, McKenna et al., 1997, Novak & Fane, 2004, Siden & Hayashi, 1974). We also find significantly reduced internal scaffolding protein B production in the decompressed ϕX174 infection (Figure 5), pointing towards another potential reason for reduced viral plaque and burst-size (Figure 2A). A lack of internal scaffolding protein B would prevent intermediary assembly particles, 9S and 6S, from forming 12S particles (Novak & Fane, 2004), thereby stalling productive virion assembly (Novak & Fane, 2004).

In the decompressed ϕX174 genome gene B has been moved from its nested position within genes A/A* to a fully separated and downstream position (Figure 6). Furthermore, the gene B 5’-UTR region is identical up to 15 nucleotides 5’ of the gene B start codon but further upstream is completely different (Figure 6) and may be a factor for its reduced protein output.

To understand possible reasons for the large differences in protein expression between decompressed and wild-type ϕX174, we analysed mRNA folding across gene 5’ termini and entire coding sequences (Appendix Figure S8) as well as host codon adaptability (Appendix Figure S9). Other work focused only on synonymous codon changes within ϕX174 had found correlations between gene mRNA folding energy and codon adaptation index (CAI) (Jaschke, Dotson et al., 2019, Van Leuven, Ederer et al., 2020). In contrast, our analyses showed that neither predictive mRNA structure or codon usage bias differences explain the difference in protein production between the strains, further emphasising our current inability to accurately predict the effects of large-scale genome architectural changes.

## Conclusion

Re-engineering and recapitulating natural genes and gene pathways of interest is now enabling researchers in synthetic biology to create genomes with novel configurations never before seen in natural systems. However, our understanding of the intricacies of genome architecture, such as the functional importance of overlapping genes, is still lacking.

In this study, we have revealed the biological ramifications of whole genome modularisation in the model virus ϕX174. For the first time, using the fully refactored ϕX174 and careful phenotypic and targeted proteomic measurements, we have conclusively demonstrated that removing coding overlap causes major defects in protein expression and reproduction efficiency. More specifically, we show that the decompressed ϕX174 infection cycle has altered protein expression for 5 of the 11 (∼45%) phage proteins (A*, B, C, E, and H), as well as perturbed burst-size, capsid stability, and host-cell attachment. This work provides key evidence that gene overlap is a crucial feature in genomes and points at a direct mechanism for their retention during evolutionary selection. Lastly, our work shows that in order to retain the highest fitness levels possible, conserving overlapped gene topologies should be strongly considered in genome engineering projects going forwards.

## MATERIALS AND METHODS

### Strains

Phage strains ϕX174 and ϕX174.1f (termed decompressed ϕX174 throughout) were used throughout this work (Jaschke et al., 2019, Jaschke et al., 2012). The host strain used for experiments and virus propagation (unless stated otherwise) was *E. coli* C122 (Public Health England NCTC122) which carried a plasmid pJ804(Gene F) containing the ϕX174 gene F (Genbank No. NC_001422.1) under a rhamnose-inducible promoter (Jaschke et al., 2012). Growth of the host strain was conducted at 37°C, 250 RPM (with 25 mm orbital shaking) in LB Miller broth (10 g/L tryptone, 10 g/L NaCl, and 5 g/L yeast extract) containing 50 μg/mL carbenicillin and 2 mM CaCl_2_ and either 2 mM rhamnose for decompressed ϕX174 or 2 mM glucose for wild-type ϕX174.

### Phage infection

Infections with either the wild-type or decompressed phage were conducted by first growing the host strain overnight (∼18-hours) followed by subculturing the cells 1/100 in fresh LB broth (Miller) containing 50 μg/mL carbenicillin and 2 mM calcium chloride, and growing until an O.D of 0.5-0.6. The cells were then pelleted (4,000 RCF, 14°C, 8-minutes), and subjected to two wash and pellet cycles with ice-cold HFB buffer (60 mM ammonium chloride, 90 mM sodium chloride, 100 mM potassium chloride, 1 mM magnesium sulfate, 1 mM calcium chloride, and 100 mM tris base, pH 7.4). After which the cell pellet was resuspended to 1/10 of the original culture volume using ice-cold HFB followed by addition of virus at required MOI. The cell-virus mixture was left at 14°C for 30-minutes to allow viral attachment (infection synchronisation) (Newbold & Sinsheimer, 1970a). Infection initiation was triggered by addition of pre-warmed (37°C) LB broth (Miller) containing 50 μg/mL carbenicillin and 2 mM CaCl_2_ (and either 2 mM rhamnose or glucose) to a final volume constituting original culture volume.

### Phage isolation and purification

Phage stocks were produced through two methods. Both involved growth of 800 mL of *E. coli* C122(pJ804(Gene F)) culture to mid-log, as described. If propagation of decompressed ϕX174 was required, the media included 50 μg/mL carbenicillin and 2 mM rhamnose. Virus was spiked into the culture and allowed to propagate until culture clearance. The first method, involved pelleting of cleared cultured to remove bacterial debris (4,000 RCF for 8-minutes). The supernatant was retained and filtered using Amicon Ultra-15 Centrifugal Filter Units (Merck, Germany) with a 100 kDa molecular weight cut-off. The second method was for ultra-pure phage stocks. After culture clearance, NaCl was added to a final concentration of 1 M, cooled to 4°C, followed by pelleting of bacterial debris (4,000 RCF for 8-minutes) and retaining the supernatant and addition of poly-ethylene glycol (PEG) 8000 to a final concentration of 15% (w/v). The supernatant was stored overnight (∼18-hours) at 4°C, followed by pelleting the phage at 4,000 RCF for 8-minutes. Phage pellets were resuspended in HFB buffer and PEG 8000 was removed using three wash cycles of equal volume chloroform. Finally, the phage was CsCl step-gradient purified using CsCl density steps of 1.30 g/mL, 1.40 g/mL, and 1.50 g/mL using the Beckman Coulter SW32Ti swinging bucket rotor at 32,000 RPM for 16-hours at 10°C. Phage bands were removed, and buffer-exchanged using the Amicon Ultra-15 Centrifugal Filter Units (Merck, Germany) with a 100 kDa molecular weight cut-off in HFB buffer. The titre for viral stocks was determined using the double-agar overlay method (Kropinski, Mazzocco et al., 2009). Soft agar was supplemented with 50 μg/mL carbenicillin and 2 mM rhamnose for enumeration of the decompressed ϕX174 phage titre.

### Genome folding prediction

Minimum free energy (MFE) secondary structure predictions for decompressed and wild-type ϕX174 were performed through the ViennaRNA websuite’s RNAfold server (Lorenz, Bernhart et al., 2011). Default settings were used, except for energy parameters which were set as DNA parameters.

### mRNA Structure Predictions

RNA folding simulations were performed on an 83-nt window around the start codon of each gene using the Nucleic Acid Package (NUPACK) web server with default settings (Zadeh, Steenberg et al., 2011), or using the entire coding sequence using mfold web server with default settings (Zuker, 2003). Lowest-energy structures were reported.

### Bacteriophage lysis curve measurements

Lysis curve for decompressed and wild-type ϕX174 was conducted in parallel 10 mL triplicate cultures, and grown and synchronised as previously described. 2 mM rhamnose was added to decompressed ϕX174 cultures to induce gene F on pJ804(Gene F), and 2 mM glucose to wild-type cultures to suppress the induction of the plasmid-borne gene F. Culture samples for O.D measurements were taken every 10-minutes for 60-minutes. PCR was performed on cultures at the end of the experiment to confirm there was no cross-contamination of phage strains (Appendix Figure S1).

### Bacteriophage attachment

Cultures were grown as described, except, lysis resistant *E. coli* C900 (Δ*slyD*) which contains a mutation conferring resistance to ϕX174 E-mediated lysis (Roof, Horne et al., 1994) was used as the host cell for attachment (a gift from Bentley A. Fane). At mid-log, cells were pelleted, washed and resuspended in microcentrifuge tubes to 1/10^th^ of the original culture volume with HFB buffer that was either pre-cooled for the 14°C assay, or pre-warmed for the 37°C assay. Phage stock was then added to the cells, followed by incubation at the required temperature with continuous culture agitation at 900 RPM using a benchtop heating block with Eppendorf holding cassette. At 5-minutes and 10-minutes post-incubation, cells were immediately pelleted at 4,000 RCF for 5-minutes, followed by removal of the top 100 µL of the supernatant for assessment of viral titre using the double-agar overlay method.

### Bacteriophage progeny production measurements

Cultures and infection synchronisation was performed as described, except, virus was added to a final MOI of 0.01. After synchronisation, cells were pelleted and the supernatant was retained for titre. Cultures were then re-suspended using pre-warmed (37°C) LB with 2 mM CaCl_2_ and 50 μg/mL carbenicillin, and supplemented with either 2 mM glucose for the wild-type virus, or 2 mM rhamnose for the decompressed phage. Resuspended cultures were then diluted with media according to pre-determined levels to allow for appropriate enumeration of plaques. Cultures were then incubated at 37°C with 250 RPM agitation for 40-minutes with sampling of the culture at 5, 10, 15, 20, 25, 30, 35, and 40-minute time-points. Sampling involved removing 200 µL of the culture and immediately mixing with 100 µL of chloroform, followed by storage on ice. Samples were then spun at 5,000 RCF for 5-minutes, after which 100 µL of the top layer was removed and put into a fresh tube. Assessment of viral titre was then performed using the double-agar overlay method.

### Virion heat stability

Phage stocks were diluted to approximately 10^2^ PFU/µL using LB broth (Miller) supplemented with 10% (v/v) glycerol, and 2 mM calcium chloride. Eppendorf tubes containing 90 µL of the same LB broth diluent as well as 10 µL of the diluted phage stock. The tubes were individually labelled according to pre-determined harvest points of 0, 5, 15, 30, and 45-minutes, with the 0-minute tube being placed immediately on ice. The rest of the tubes were incubated at 60°C, and then placed upon ice when reaching their time point. Titres of the samples was performed as described using the double-agar overlay method.

### DNase I and proteinase K stability assays

For both assays, individual plaques were stabbed from double-agar overlay plates and eluted into HFB buffer for the DNase I assay, or 10 mM Tris-HCl for the proteinase K assay. The DNase I assay consisted of aliquoting 7.5 μL of decompressed or wild-type plaque suspension into a 200 μL capacity microcentrifuge tubes containing 0.5 μL DNase I (2,000 U/mL) (or water (control)), 1 μL DNase I buffer (10x) and incubated at 37°C for 1-hour. The proteinase K assay consisted of aliquoting 9 μL of plaque suspension into a 200 μL capacity microcentrifuge tube and either 1 μL of 1 μg/μL proteinase K, or water (control) and incubated at 37°C of for 4-hours. For both assays, plaques were enumerated through the double-agar overlay assay (Kropinski et al., 2009).

### Electron microscopy

A 4 μL solution of decompressed or wild-type ϕX174 virions were applied to glow-discharged copper grids (carbon type B with formvar, 200 mesh (Ted Pella Inc., United States)), then stained with 2% uranyl acetate and dried at room temperature. The screening of the capsids was carried on FEI Tecnai G2 20 electron microscope (ThermoFisher Scientific, United States) at 200 kV accelerating voltage. Data images for 2D classification of both capsids were automatically collected on a Talos Arctica transmission electron microscope operated at 200 kV acceleration voltage using EPU software (ThermoFisher Scientific, United States). All datasets were collected on a Falcon III detector in linear mode at 92,000× magnification with a pixel size of 1.6 Å per pixel.

### Image processing (2D classification)

MotionCor2 (Zheng, Palovcak et al., 2017) was used to correct local beam-induced motion and to align resulting frames. Micrographs were then sorted on the basis of image quality, and 1,946 micrographs for wild-type & 1,633 micrographs for decompressed ϕX174 were used for subsequent reconstruction using RELION 3.0 beta (Fernandez-Leiro & Scheres, 2017, Scheres, 2012b, Zivanov, Nakane et al., 2018). Particles (37,150 for wild-type & 2,971 for decompressed ϕX174) were picked automatically from the full data set and extracted by using a box size of 300 × 300 pixels. The particles were then analysed by 2D classification in RELION 3.0 beta.

### Cryo-electron microscopy

A 4 μL solution of co-isolated small particles were applied to glow-discharged copper grids (Quantifoil R2/2, Quantifoil Micro Tools), blotting for 3-seconds at 4 °C with 90% humidity and plunging into liquid ethane using a Lecia EM GP device (Leica Microsystems, Germany). Cryo-EM data were collected on a Talos Arctica transmission electron microscope operated at 200 kV acceleration voltage using EPU automated data collection software (ThermoFisher Scientific, United States). All datasets were collected on a Falcon III detector in counting mode at 120,000× magnification with a pixel size of 1.25 Å per pixel.

### Image processing and refinement

MotionCor2 (Zheng et al., 2017) was used to correct local beam-induced motion and to align resulting frames. Defocus and astigmatism values were estimated using Gctf (Zhang, 2016). Micrographs were then sorted on the basis of image quality, and 1,424 micrographs were used for subsequent reconstruction using RELION 3.0 beta (Fernandez-Leiro & Scheres, 2017, Scheres, 2012b, Zivanov et al., 2018). ∼1,000 particles from the data set were manually picked and subjected to 2D classification to generate templates for auto-picking. 72,212 particles were automatically picked and extracted by using a box size of 200 × 200 pixels. The particles were analysed by 2D classification, and 10 classes showing clear structural details were selected for further processing with 61,922 particles. The map at 20 Å generated from Protein Data Bank (2C7E) was used as an initial model for 3D classification. A subset of 23,517 particles was then used for the first round of 3D refinement, which results in a map with a resolution of 5.7 Å. The resolution was improved to 5.1 Å after doing post-processing (soft-edge masking and B-factor sharpening). 3D maps were post-processed to automatically estimate and apply the B-factor and to determine the resolution by Fourier shell correlation (FSC) between two independent half datasets using 0.143 criterion (Scheres, 2012a).

### Label-free quantitative mass spectrometry – Parallel reaction monitoring (PRM)

*E. coli* C122(pJ804(Gene F)) was grown from overnight (∼18-hours) cultures to mid-log phase in LB broth (Miller) supplemented with 2 mM CaCl_2_ and 50 μg/mL carbenicillin. Cultures were then prepared for infection synchronisation with either decompressed or wild-type ϕX174 strains at MOI = 1.2 (to the starved hosts, we added 230 μL of LB broth (Miller) with 2 mM CaCl_2_ and 50 μg/mL carbenicillin containing phage at the appropriate PFU), as described earlier. After which, the infection was initiated by the addition of warm (37°C) LB broth (Miller) supplemented with 2 mM CaCl_2_, 50 μg/mL carbenicillin and either 2 mM rhamnose for the decompressed ϕX174 strain cultures, or glucose for the wild-type cell cultures. 0-minute time-points were collected immediately (2.25 mL) and placed on ice and then the cultures were incubated at 37°C, 250 RPM. At time-points 15, 25, and 35-minutes post-infection initiation, 2.25 mL of culture was harvested and stored on ice. Samples were pelleted, and washed twice with ice-cold HFB buffer, followed by storing the pellets at -30°C overnight (∼18-hours).

The next day, sample pellets were resuspended in 200 µL lysis buffer (100 mM triethylammonium bicarbonate buffer (TEAB) with 1% (v/v) sodium deoxycholate (SDC) and 1x protease inhibitor cocktail solution (Roche, Switzerland)), and subjected to probe sonication (30% amplitude, 5 bursts). Reduction and alkylation of cysteine residues (dithiothreitol (DTT) to 10 mM, incubated at 60°C for 30-minutes followed by iodoacetamide (IAA) to 30 mM, incubated 1-hour in the dark, and lastly, quenching with equal molarity of DTT), followed by chloroform-methanol protein precipitation to obtain pure protein pellets that were subsequently resuspended in digestion buffer (100 mM TEAB with 1% (v/v) SDC) and quantified by the Pierce™ BCA Protein Assay Kit (ThermoFisher Scientific, United States). From each sample, 30 μg of protein was taken for digestion with trypsin (2% (w/w)) (Promega, United States) and left overnight at 37°C (∼18-hours). After digestion, samples were acidified with 100% formic acid to 1% (v/v) and SDC was removed by spinning samples at 20,000 RCF and collecting the supernatant into a fresh tube. Samples were then vacuum centrifuged and dry peptides were stored at -30°C until ready for mass spectrometry analysis.

Peptides were reconstituted in 2% (v/v) acetonitrile (ACN) and 0.1% (v/v) formic acid (buffer A) to a final concentration of 0.1 μg/µL. 1μg (10 μL) of each sample was analysed on a Q-Exactive mass spectrometer (ThermoFisher Scientific, United States) coupled to an EASY-nLC1000 system (ThermoFisher Scientific, United States). Peptide samples were injected onto the LC system using buffer A and were bound on a 75 µM x 100 mM C18 HALO column (2.7 µM bead size, 160 Å pore size). A flow rate of 300 nL/min using an increasing linear gradient of buffer B (99.9% (v/v) acetonitrile, 0.1% (v/v) formic acid) was run from 1% to 50% for 110-minutes followed by 85% buffer B for 10-minutes.

An inclusion list was used to target pre-defined precursor *mass-to-charge* (*m/z*) (Appendix Dataset S3), and the mass spectrometer was operated to perform one full-ms scan (70000 resolution (at 400 *m/z*) across the *m/z* range of 320-1800 *m/z*) with an automatic gains control (AGC) target of 1e6 (or a maximum fill time of 200 ms), followed by sequential PRM scans using the inclusion list (loop count 39, isolation window of 2 *m/z*). The selected precursor ions from PRM scans had an AGC target of 1e6 and a maximum fill time of 58 ms, after which, they were transferred from the C-trap to the higher energy collision dissociation (HCD) cell for fragmentation at a normalized collision energy of 27. MS/MS spectra were collected at a resolution of 17500 (at 400 *m/z*).

Raw files were analysed by MaxQuant version 1.6.10.43 (Cox & Mann, 2008) against a ϕX174 protein sequence list generated from GenBank file NC_001422.1. Cysteine carbamidomethylation was selected as a fixed modification and protein and peptide sequence match identifications were set at a 1% false discovery rate cut-off. Result files were imported and analysed in Skyline version 19.1.0.193 (MacLean, Tomazela et al., 2010), and peptides with less than 3 matching fragment ions were filtered out. The summed area values of all individual transitions for each precursor (total area) were extracted after applying Savitzky-Golay Smoothing and these values were used to quantify peptide relative abundance. The median total ion current (TIC) area across all samples was used for normalisation by correcting for injection amount. The median TIC was divided by the sample’s TIC and the ratio of this was used to normalise the total transition area. Average and standard deviation of the total area were calculated across biological triplicates, constituting relative peptide (and by extension, protein) abundance. Statistical analysis was performed individually on each peptide and significance level was established using single or two-factor ANOVA where appropriate (significance = p-value < 0.05). Fold-change was expressed using decompressed ϕX174 average intensity over wild-type ϕX174 average intensity.

## ACKNOWLEDGEMENTS

We recognize that this research was conducted on the traditional lands of the Wattamattagal clan of the Darug nation. BWW was supported by a Macquarie Research Excellence PhD Scholarship, PRJ was supported by the Molecular Sciences Department, Faculty of Science & Engineering, and the Deputy Vice-Chancellor (Research) of Macquarie University. JFR thanks the Cryo Electron Microscopy Facility through the Victor Chang Innovation Centre, funded by the NSW government, and the Electron Microscope Unit at UNSW, Sydney. We thank Dr. Pascal Steffen (The University of Sydney), Dr Roy Walker (Macquarie University), and Prof. Bentley Fane (University of Arizona) for helpful discussions and feedback, Dr. Chao Shen (Macquarie University) for assistance with microscopy. Aspects of this research were conducted at the Australian Proteome Analysis Facility.

## AUTHOR CONTRIBUTIONS

BWW contributed Conceptualisation, Formal analysis, Investigation, Methodology, Visualisation, Writing - original draft, and Writing - review and editing. JR contributed Formal analysis, Investigation, Methodology, Resources, Validation, Visualization, Writing – original draft, and Writing – review & editing. PRJ contributed Conceptualization, Funding acquisition, Project administration, Resources, Validation, Visualization, Writing - review & editing, and Supervision. MPM contributed Conceptualization, Funding acquisition, Resources, Writing - review & editing, Supervision.

## DATA ACCESS

Raw and processed proteomics data generated in this study is available in the Panorama Repository under https://panoramaweb.org/phix174.url and ProteomeXchange under PXD019681.

