## Appendix for "Genome modularization reveals overlapped gene topology is necessary for efficient viral reproduction"

**CONTENTS**

APPENDIX FIGURES

Figure S1: Representative sample purity test using PCR.

Figure S2: Computational prediction of 2D folding structure of wild-type and decompressed genomes.

Figure S3: Phage stability assays for decompressed (D) and wild-type (WT).

Figure S4: Co-isolated small particles revealed as likely *E. coli* GroEL chaperone complex

Figure S5: Decompressed and wild-type φX174 capsid 2D class averaging

Figure S6: Summed average transition ion intensity of peptides as measured across four time-points. Proteins A-E.

Figure S7: Summed average transition ion intensity of peptides as measured across four time-points. Proteins F-K.

Figure S8: Predicted N-terminal mRNA lowest energy structure is not predictive of phage protein production.

APPENDIX TABLES

Table S1: Genes of φX174

APPENDIX FILES

File S1: Peptides and final results.

File S2: PRM results for viral peptides.

File S3: PRM inclusion list

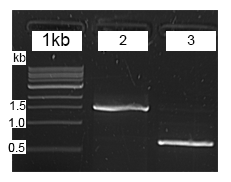

**Figure S1:** **Representative sample purity test using PCR.** Electrophoresis gel as quality control to ensure no strain cross-contamination during the lysis curve measurements (Figure 1B). Lane 1: 1 kb ladder, lane 2: wild-type growth curve sample, lane 3: decompressed φX174 growth curve sample. The gel indicates no detectable contamination between samples.

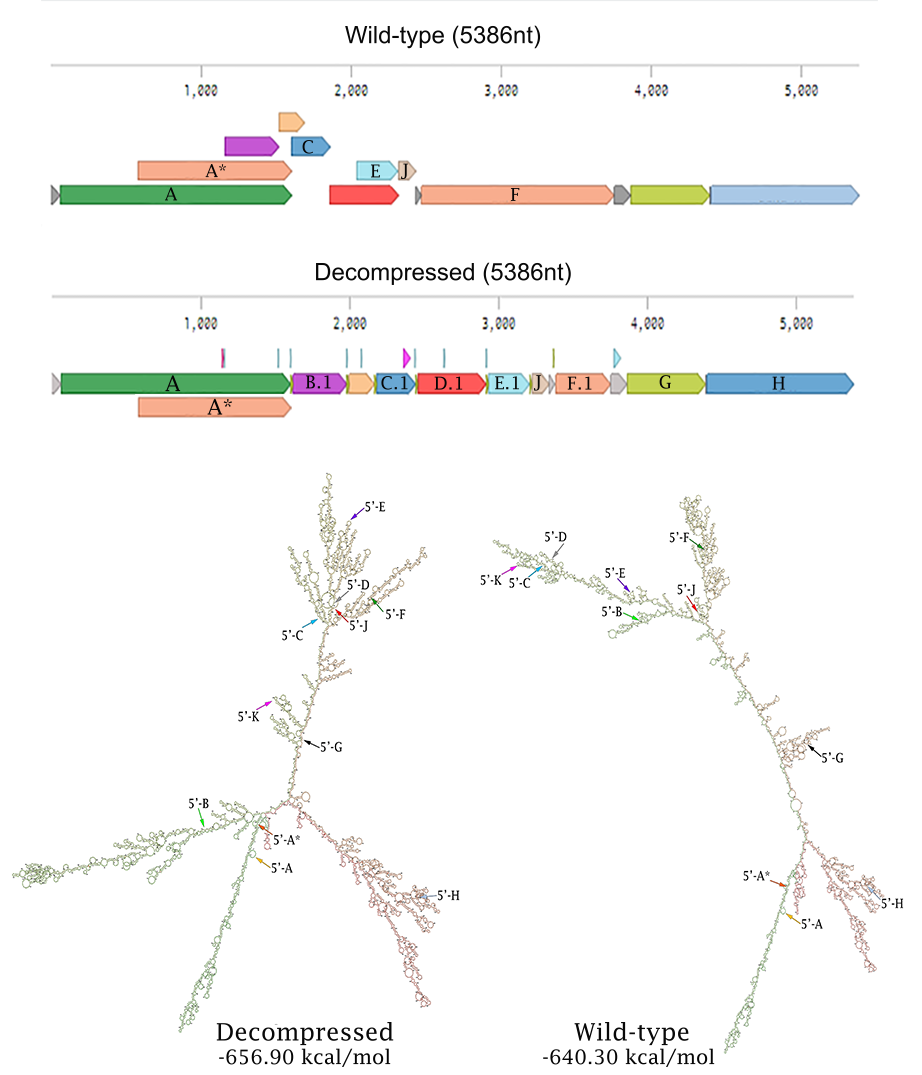

**Figure S2:** **Computational prediction of 2D folding structure of wild-type and decompressed genomes.** Linearised genomes (top) and the RNAfold predicted minimum free energy secondary structures of the φX174 genomes (bottom) as computed through RNAfold server [1]. Gene names are annotated on the linearised genomes, and 5’- gene start sites have been annotated on the structure plots.

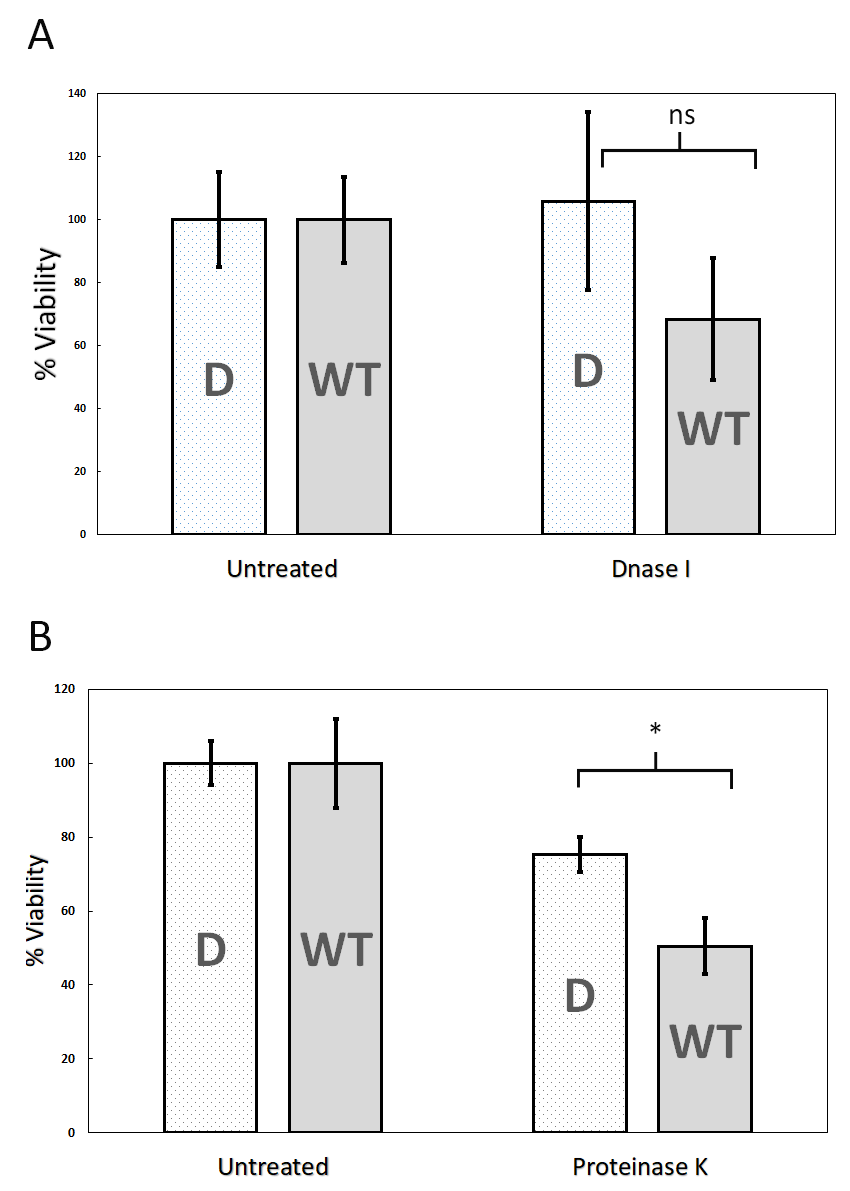

**Figure S3: Phage stability assays for decompressed (D) and wild-type (WT).** (A) Purified phage was untreated or incubated with DNase I for 1-hour at 37°C (n = 4), after which viability was determined by plaque enumeration. (B) Purified phage was untreated or incubated with proteinase K for 4-hours at 37°C (n = 3), after which viability was determined by plaque enumeration. Error bars represent standard deviation. ns = no-significance and * = p-value 0.0173.

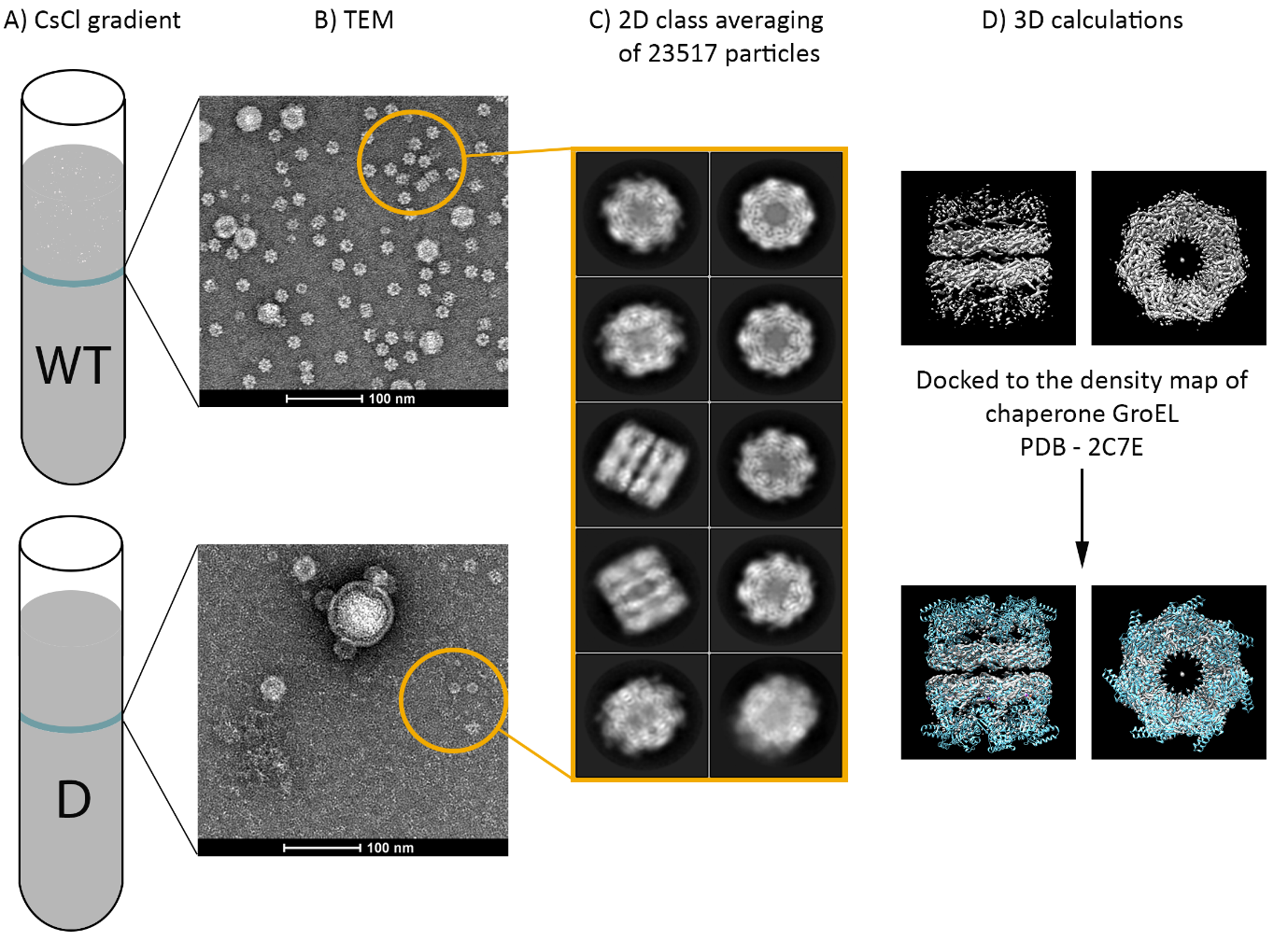

**Figure S4: Small particles co-isolated with φX174 virions revealed as likely *E. coli* GroEL chaperone complex.** (A) Phage band isolated after CsCl step-gradient ultracentrifucation. WT = wild-type and D = decompressed. (B) Uranyl-acetate negative stained particles visualised using transmission electron microscopy (TEM). Smaller particles found in both preparations (orange circle). (C) 2D class averaging of smaller particles highlighted in (B). (D) 3D representation of small particles reveal structural information and good fit to *E. coli* chaperone complex GroEL.

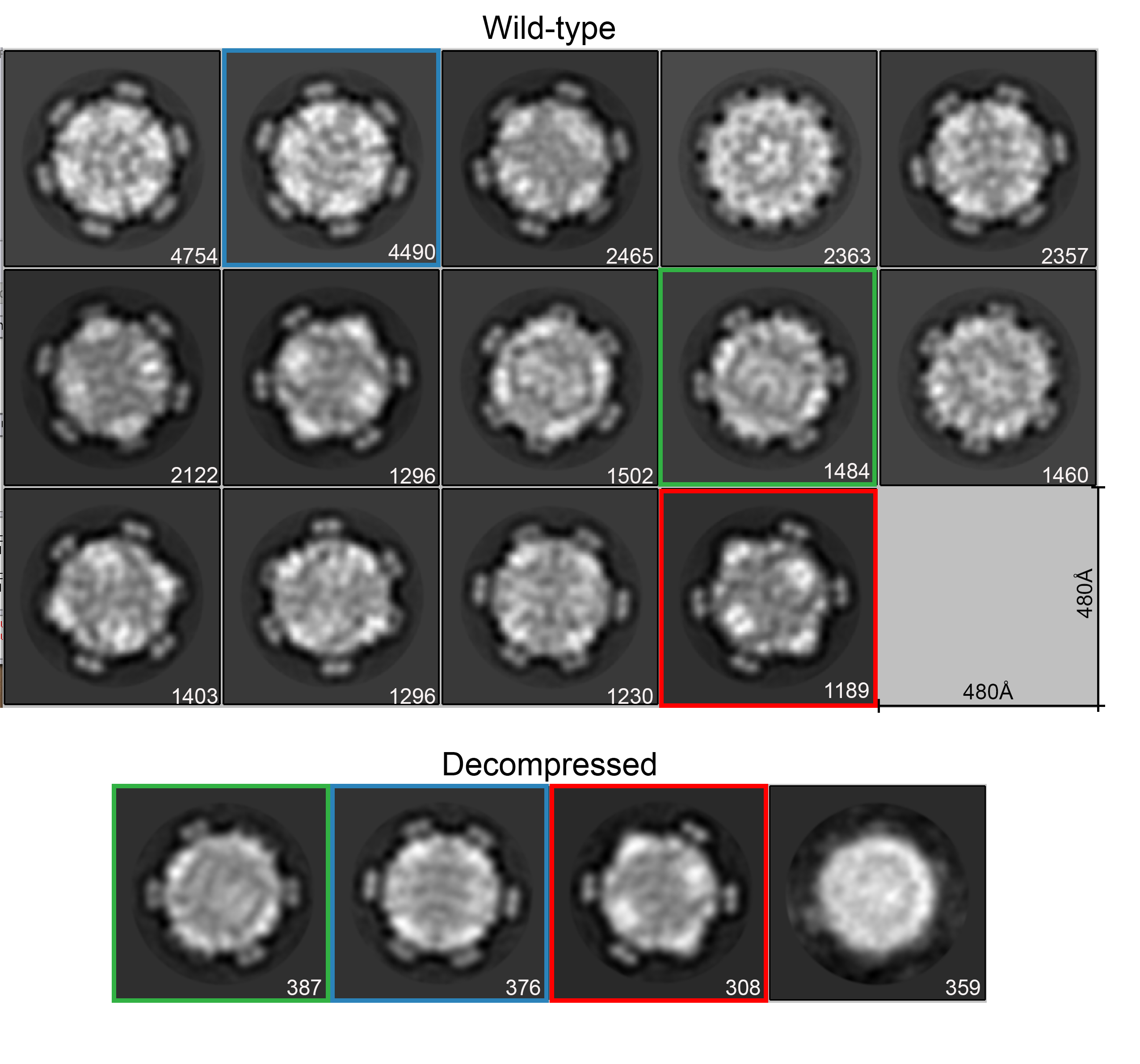

**Figure S5: Decompressed and wild-type φX174 capsid 2D class averaging.** Uranyl-acetate negative stained particles visualised using transmission electron microscopy followed by 2D class averaging. Values in bottom right corner of each image represent number of particles used for averaging. Coloured boxes indicate shared class between wild-type and decompressed φX174.

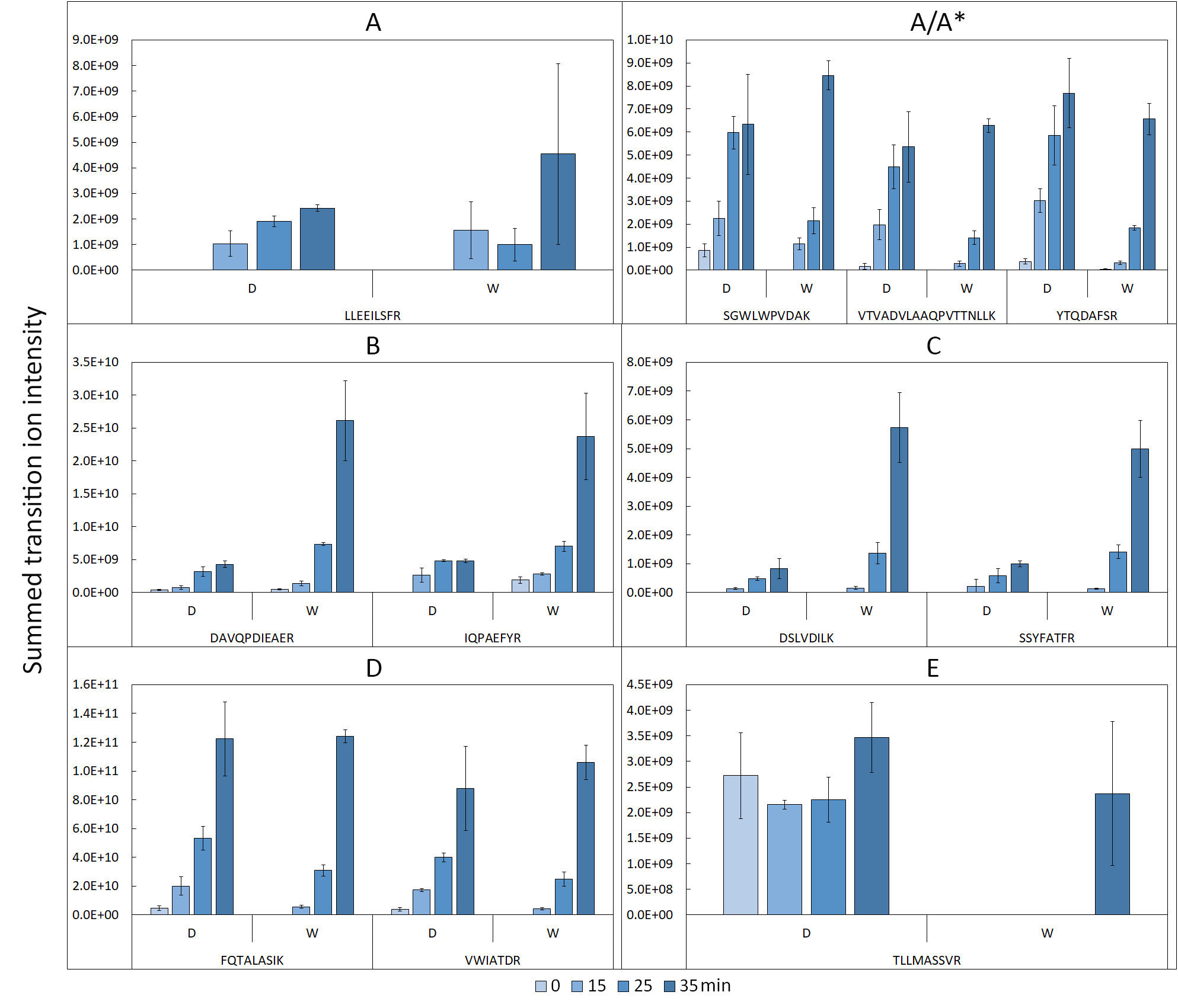

**Figure S6: Summed average transition ion intensity of peptides from proteins A/A*, B, C, D, E across infection cycle.** Each plot depicts the relative abundance of targeted peptides. Error bars respresent one standard deviation (n = 3). D = decompressed strain, W = wild-type strain. Accompanying statistical analysis in Supplementary File S1 and S2.

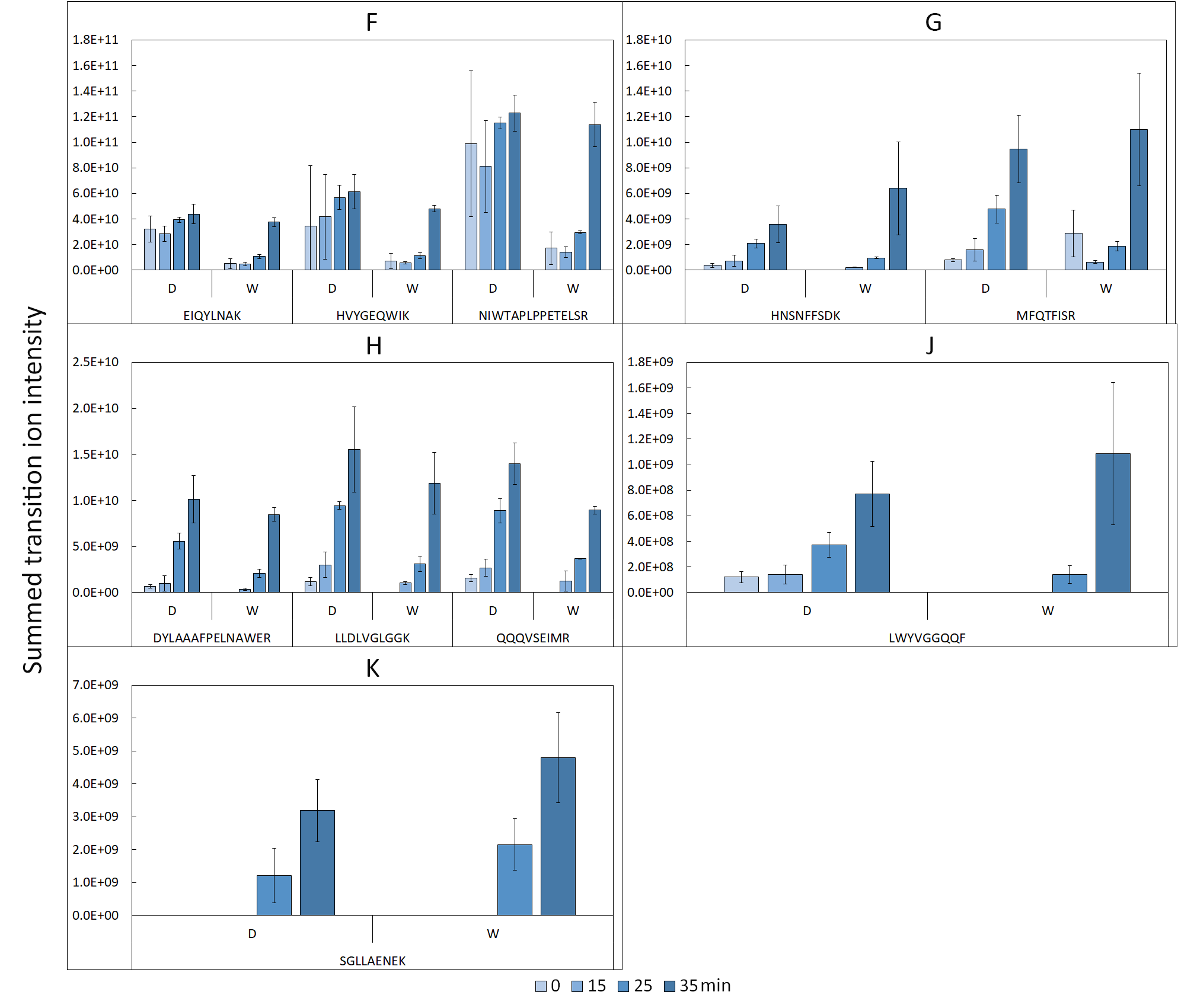

**Figure S7: Summed average transition ion intensity of peptides from proteins F, G, H, J, K across infection cycle.** Each plot depicts the relative abundance of targeted peptides. Error bars respresent one standard deviation (n = 3). D = decompressed strain, W = wild-type strain. Accompanying statistical analysis in Supplementary File S1 and S2.

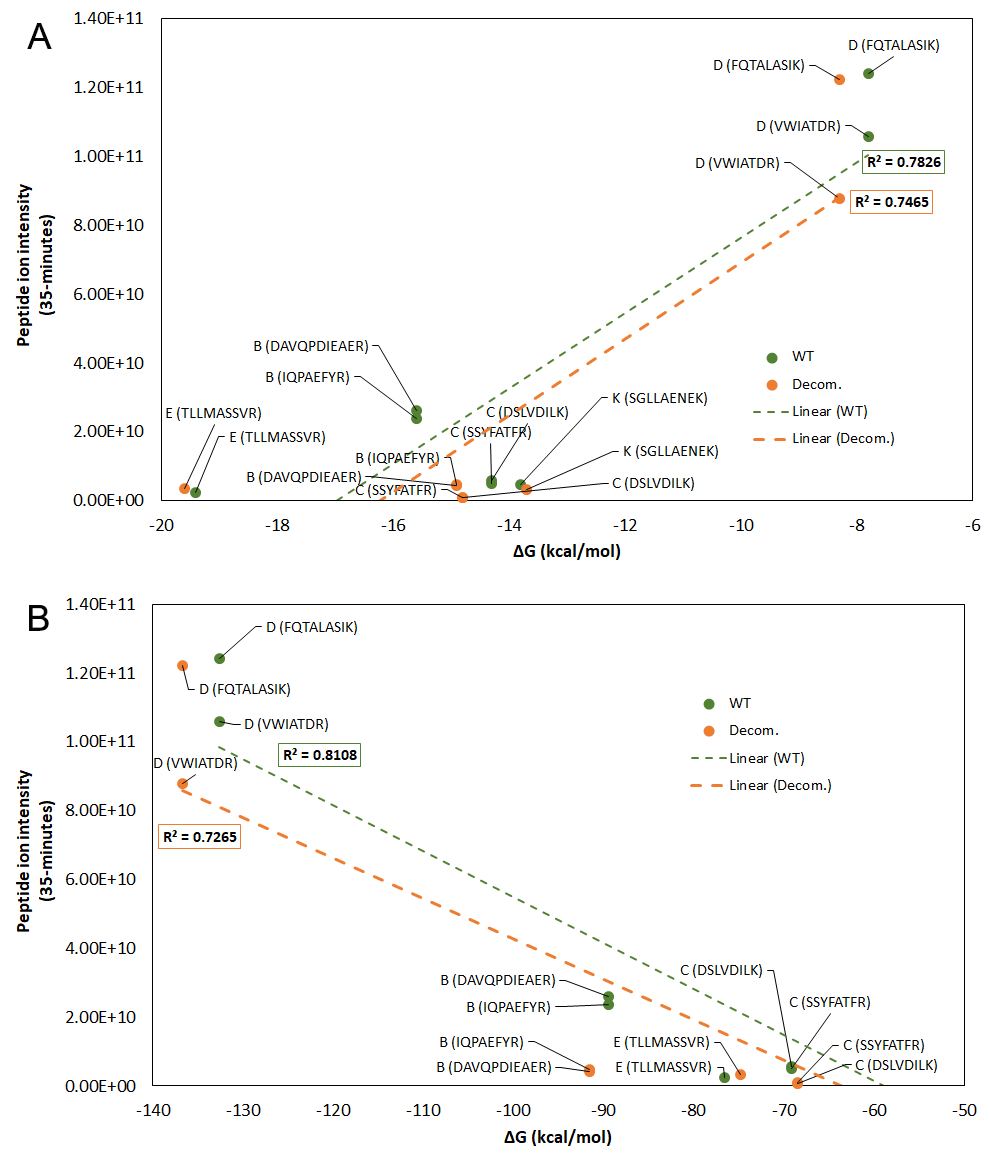

**Figure S8**: **Predicted mRNA lowest energy structure is not predictive of phage protein production differences between the strains**. For φX174 genes that were largely modularized during the refactoring process (Supplementary figure S2) and therefore mRNA structure likely impacted, RNA folding simulations were performed on each gene and resulting lowest energy structures plotted against relative peptide intensity (35-minutes post-infection). (A) 83-nt windows around the start codon of each ORF was used for the calculation using NUPACK (default settings) [2], and (B) the entire coding sequence was used as the input into mfold (default settings) [3] for energy calculations. Lowest energy structures reported. Peptide identities reported within brackets.

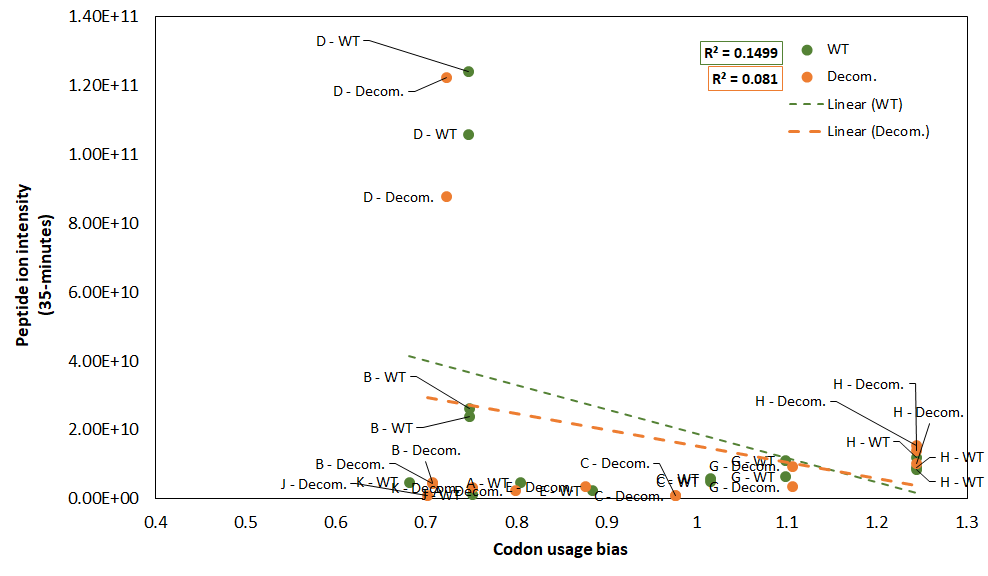

**Figure S9: Codon usage bias is not predictive of protein production differences between strains.** Codon bias discrepancy between φX174 genes and host genome codon adaptability index was calculated for decompressed (decom.) and wild-type (WT) using the MILC method [4]. Resultant codon usage bias for each gene was plotted against relative peptide intensity (35-minutes post-infection). Genes and strain are reported in callout boxes.

**Table S1. Genes of φX174.**

| **Gene** | **Gene product function** | **Reference(s)** |
| --- | --- | --- |
| **A** | φX174 DNA replication (stages II and III) through binding to the origin of replication. | Francke and Ray [5], Eisenberg, et al. [6], Fujisawa and Hayashi [7], Eisenberg and Kornberg [8] |
| **A*** | Regulates φX174 DNA replication through competitive binding of the stage II DNA thereby preventing the activity of A and encouraging stage III synthesis. Inhibits host DNA replication. Not essential. | Eisenberg and Ascarelli [9], Colasanti and Denhardt [10], van Mansfeld, et al. [11], Colasanti and Denhardt [12], Roznowski, et al. [13] |
| **B** | Internal procapsid scaffolding.  (60 copies) | Mukai, et al. [14], Dokland, et al. [15] |
| **C** | DNA replication stage II to III DNA synthesis switching through competitive binding of the origin of replication. Packaging of the ssDNA genome in conjunction with protein J. | Aoyama and Hayashi [16], Doore, et al. [17] |
| **D** | External procapsid scaffolding.  (240 copies) | Fujisawa and Hayashi [18], Dokland, et al. [15] |
| **E** | Cell lysis by inhibition of host peptidoglycan synthesis. | Hutchison and Sinsheimer [19], Pollock, et al. [20], Witte, et al. [21], Bernhardt, et al. [22] |
| **F** | Major coat protein.  (60 copies) | Fujisawa and Hayashi [18], McKenna, et al. [23] |
| **G** | Major spike protein.  (60 copies) | Fujisawa and Hayashi [18], McKenna, et al. [23], |
| **H** | DNA pilot protein, forms a tail for genome transport. Phage adsorption protein by recognition of host lipopolysaccharide receptor. Guides phage DNA through host cell wall.  (12 copies) | Jazwinski, et al. [24], McKenna, et al. [23], Sun, et al. [25] |
| **J** | Mediates DNA packaging by complexing to stage II and III DNA and accompanying it into the procapsid. Tethers phage genome to the inner surface of the capsid wall (major coat protein).  (60 copies in procapsid/mature virion) | Hamatake, et al. [26], McKenna, et al. [23], Hafenstein, et al. [27], Bernal, et al. [28] |
| **K** | Burst size modulation. Not essential. | Gillam, et al. [29] |
